# Population structure and transmission of *Mycobacterium bovis* in Ethiopia

**DOI:** 10.1101/2020.11.17.386748

**Authors:** Gizat Almaw, Getnet Abie Mekonnen, Adane Mihret, Abraham Aseffa, Hawult Taye, Andrew JK Conlan, Balako Gumi, Aboma Zewude, Abde Aliy, Mekdes Tamiru, Abebe Olani, Matios Lakew, Melaku Sombo, Solomon Gebre, Colette Diguimbaye, Markus Hilty, Adama Fané, Borna Müller, R Glyn Hewinson, Richard J Ellis, Javier Nunez-Garcia, Eleftheria Palkopoulou, the ETHICOBOTS consortium, Tamrat Abebe, Gobena Ameni, Julian Parkhill, James LN Wood, Stefan Berg, Andries J van Tonder

## Abstract

Bovine tuberculosis (bTB) is endemic in cattle in Ethiopia, a country that hosts the largest national cattle herd in Africa. The intensive dairy sector, most of which is peri-urban, has the highest prevalence of disease. Previous studies in Ethiopia have demonstrated that the main cause is *Mycobacterium bovis* (*M. bovis*), which has been investigated using conventional molecular tools including deletion typing, spoligotyping and Mycobacterial interspersed repetitive unit-variable number tandem repeat (MIRU-VNTR). Here we use whole genome sequencing (WGS) to examine the population structure of *M. bovis* in Ethiopia. A total of 134 *M. bovis* isolates were sequenced including 128 genomes from 85 mainly dairy cattle and six genomes isolated from humans, originating from 12 study sites across Ethiopia. These genomes provided a good representation of the previously described population structure of *M. bovis*, based on spoligotyping and demonstrated that the population is dominated by the clonal complexes African 2 (Af2) and European 3 (Eu3). A range of within-host diversity was observed amongst the isolates and evidence was found for both short- and long-distance transmission. Detailed analysis of available genomes from the Eu3 clonal complex combined with previously published genomes revealed two distinct introductions of this clonal complex into Ethiopia between 1950 and 1987, likely from Europe. This work is important to help better understand bTB transmission in cattle in Ethiopia and can potentially inform national strategies for bTB control in Ethiopia and beyond.

## Introduction

*Mycobacterium bovis* (*M. bovis*) is one of several highly related subspecies of the *Mycobacterium tuberculosis* complex (MTBC) which causes tuberculosis (TB) in humans and a range of domesticated and wild animals [1]. While the main human pathogen of this complex, *Mycobacterium tuberculosis* sensu stricto, causes approximately ten million new TB cases yearly [2], *M. bovis* is the main causative agent of TB in cattle, also known as bovine TB (bTB). The actual global prevalence of bTB in cattle is not known but recent estimates suggest that around 7% of all cattle around the world are likely affected, having a significant impact on their productivity [3]. *M. bovis* is also capable of being transmitted to humans and it is believed that at least 1.5% of all human TB cases are due to zoonotic TB [2].

Bovine TB is present across all continents and many countries in Europe, Australasia and the Americas have been able to eliminate or control bovine TB in their national herds, often through costly test-and-slaughter programmes [4, 5]. In countries without adequate resources to control the disease, bTB is frequently endemic [2]. This is the case for Ethiopia which has the fifth largest national cattle herd in the world with over 60 million animals, of which the vast majority are local zebu breeds reared in extensive farming systems [6]. bTB endemic at a very low level in these extensive husbandry systems but thrives in the intensive dairy sector, with mainly high milk-yield Holstein-Friesian (H-F) or H-F/zebu cross-bred dairy cattle, where the prevalence is high [7]. This is particularly true in the well-established dairy belt in central Ethiopia where several studies have recorded over 25% bTB prevalence in animals [8, 9] (Almaw *et. al*, Submitted for publication). With the current increase in urbanization in Ethiopia, demand for milk is increasing around urban centres [10] and consequently, the intensive dairy sector is expanding and emerging in new centres across the country. Such expansion is associated with trading of dairy cattle between herds and regions, leading to increased risk of disease transmission in this sector.

To address questions around transmission and control of bTB in the highly affected diary sector, a project named Ethiopia Control of Bovine TB Strategies (ETHICOBOTS) was funded by the Zoonoses and Emerging Livestock Systems programme (UK). One aim of this multi-disciplinary project was to investigate the population structure of *M. bovis* in the Ethiopia dairy sector and to explore epidemiological links. Previous studies have used strains from across Ethiopia to establish a basic picture of the *M. bovis* population through conventional methods including spoligotyping and Mycobacterial interspersed repetitive unit-variable number tandem repeat (MIRU-VNTR) [8, 11–19]; this contributed to the definition of the African 2 clonal complex of *M. bovis* (Af2) confined to Ethiopia and East Africa [20]. However, these techniques have their limitations: MIRU-VNTR has less reproducibility and spoligotyping has less discriminatory power [21–23]. Whole genome sequencing (WGS) and single nucleotide polymorphism (SNP)-based analyses can now provide more detailed and discriminating phylogenetic analyses. Also, due to a lack of recombination in the MTBC, SNPs exhibit very low degrees of homoplasy [22, 24].

A small number of *M.bovis* isolates (n = 14) from neighbouring Eritrea have previously been analysed using WGS [25]. In this study we performed genomic analyses of 134 Ethiopian *M. bovis* isolates collected from dairy cattle, humans and a dromedary from several study sites across the country, giving the first WGS-based picture of the population structure of *M. bovis* in Ethiopia.

## Methods

### Isolate selection, culturing, and preparation of genomic DNA

Metadata on the host species of the 134 *M. bovis* isolates selected for this study are summarized in Supplementary File 1. The vast majority (n = 121) originated from 76 dairy cattle (12 H-F, 8 Zebu and 56 H-F/Zebu cross-bred dairy cattle), collected from known dairy farms in Mekele, Gondar, and Alage (near Hawassa) as well as from the large dairy belt in central Ethiopia (Addis Ababa, Holeta, Sebeta, Bishoftu, Sendafa and Sululta) while six isolates were collected from zebu cattle and one from a dromedary slaughtered at Gondar, Addis Ababa, Butajira, Negele, and Filtu abattoirs respectively (Figure 1A; Supplementary File 1). The latter six, the dromedary sample, plus 29 isolates from dairy cattle from central Ethiopia were collected in previous studies [8, 11, 14] and frozen stocks (archived at −80°C) were used for re-culturing and subsequent genomic DNA preparation. Ninety-two of the isolates from dairy cattle were collected as part of the ETHICOBOTS project. Eighty-six of these isolates originated from tuberculin reactor animals that were slaughtered and tissue samples of suspected TB lesions were processed and cultured for mycobacteria according to a previously described protocol [26]. The remaining six isolates originated from milk collected from tuberculin reactors. These milk samples were processed based on published methodology [27–29] and cultured as described for the tissue samples. The six *M. bovis* isolates of human origin were isolated from pulmonary TB cases (sputum) or from TB lymphadenitis cases (fine-needle aspirates), from study sites overlapping with the cattle samples (Supplementary File 1) [30, 31] (Hawult Taye, manuscript in preparation). All *M.bovis* isolates included in this study were isolated between 2006 and 2018 (Supplementary File 1). Genomic DNA of each isolate was extracted from heat-inactivated *M. bovis* cells using the DNeasy Blood & Tissue Kit (Qiagen). Before submission for WGS, all genomic DNA samples were confirmed as *M. bovis* by RD4 deletion typing [11].

**Figure 1.**
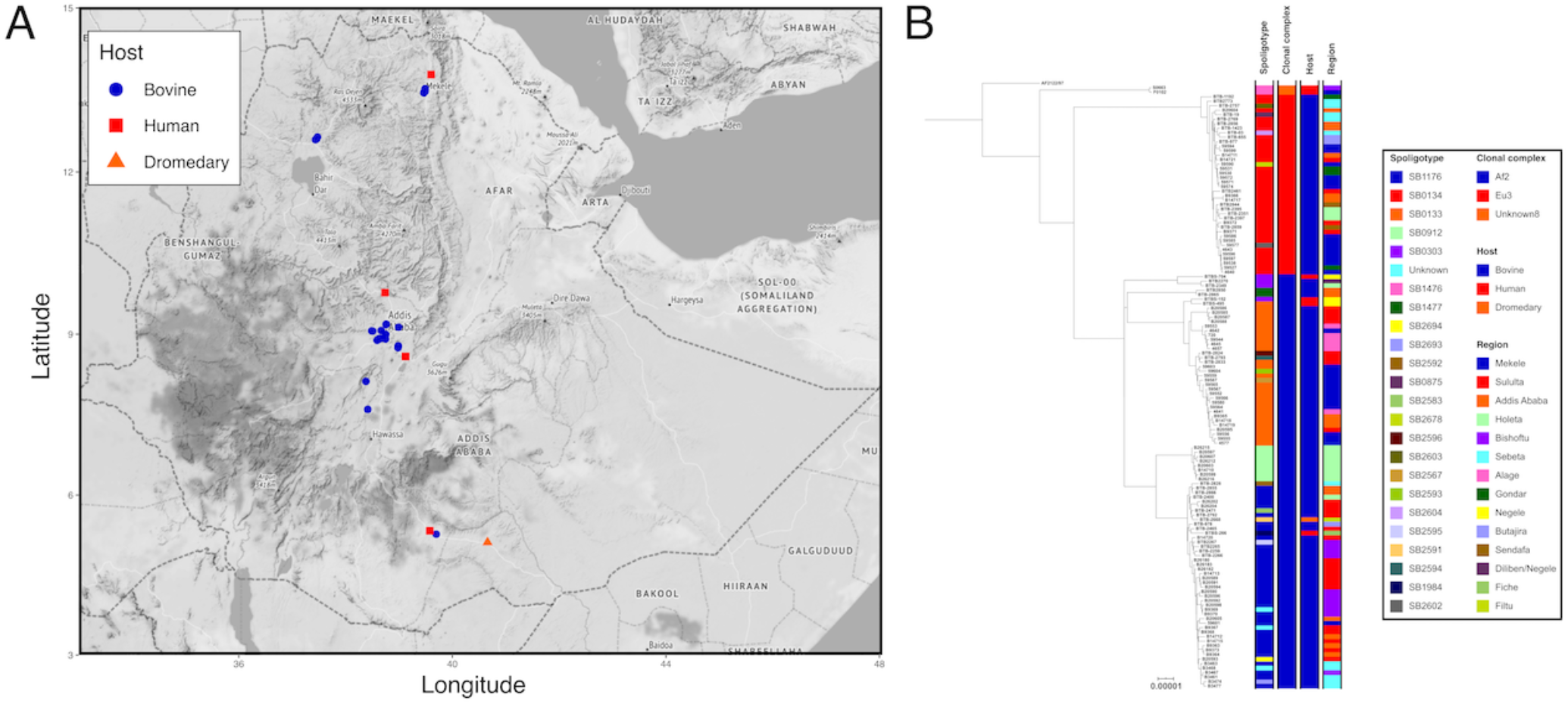
A. Map of Ethiopia showing the location of isolation for each sequenced isolate coloured by the host; B. Maximum likelihood phylogenetic tree of 134 Ethiopian genomes rooted using *Mycobacterium bovis* AF2122/97.

### Whole genome sequencing and sequence analysis

WGS was performed in four rounds using the Illumina HiSeq 2500, MiSeq, NextSeq 500, and HiSeqX platforms to produce paired-end reads of between 50 and 150 base-pairs in length. Raw sequencing reads were deposited at the European Nucleotide Archive (ENA) under project PRJEB32192; all accessions used in this project are listed in Supplementary File 1. FastQC v0.11.9 [32] was used to generate basic quality control metrics for the raw sequence data. Sequence reads were classified using Kraken v0.10.6 [33] and the abundance of the classification was refined to a single level using Bracken v1.0 [34]. Samples with less than 70% of all reads assigned to the *Mycobacterium* genus were excluded. *In silico* spoligotyping was performed using SpoTyping v2.1 [35] and the binary spoligotype representations were queried against the *M. bovis* spoligotype database (www.mbovis.org) to extract spoligotypes named in the format of SBXXXX. Clonal complexes were assigned to samples using RD-analyzer v1.0 [36] with samples not identified as belonging to previously described *M. bovis* clonal complexes (European 1 (Eu1), European 2 (Eu2), African 1 (Af1), and African 2 (Af2)) designated as “Other” [20, 37–39]. Further assignment to clonal complex was based on the phylogenetic lineages recently identified by Loiseau *et al*. [40].

Sequence reads were mapped to the *M. bovis* AF2122/97 reference genome (NC0002945) using BWA v0.7.17 (Burrow-Wheeler Aligner) (minimum and maximum insert sizes of 50 and 1000, respectively) [41]. Single nucleotide polymorphisms (SNPs) were called using SAMtools v1.2 mpileup and BCFtools v1.2 (minimum base call quality of 50 and minimum root squared mapping quality of 30) as previously described [41, 42]. Samples with reads mapping to less than 90% of the AF2122/97 reference were excluded. Genomic regions consisting of GC-rich sequences such as PPE proteins and PE-PGRS repeats were masked in the resulting alignment using previously published coordinates [43]. Variable sites were extracted from the masked alignment using snp-sites v2.5.1 [44]. Maximum likelihood phylogenetic trees were constructed using IQ-tree v1.6.5 [45] (constant sites added to alignment, extended model selection and 1000 bootstraps) and the resulting trees were annotated and rooted in iTOL [46]. Pairwise SNP distances for all genomes were calculated using pairsnp (https://github.com/gtonkinhill/pairsnp).

To provide wider context for the isolates sequenced in this study, a global representative *M. bovis* dataset (n = 352; Supplementary File 2) was assembled. Genomes were selected based on the following criteria: they were previously published [25, 40, 47–51] and represented the currently understood *M. bovis* diversity. All previously characterized clonal complexes (Af1, Af2, Eu1, Eu2, and Eu3) and uncharacterized lineages (Unknown1 to Unknown8) from Loiseau et al. [40] were included. Due to the large number of available sequences for clonal complexes Eu1 and Eu2, a random selection of 100 genomes was chosen for each. Sequence data was downloaded from the ENA and trimmed using Trimmomatic v0.33 [52]. Sample QC, spoligotype assignment, mapping and phylogenetic tree construction were performed as above. The tree was rooted with a *Mycobacterium caprae* isolate.

### Spatial analysis

A map of the geographical locations of isolate collection (latitude and longitude) was constructed using R and the ggmap library [53, 54]. The spatial position of genomes from each clonal complex was plotted and a convex hull (the smallest polygon incorporating a given set of points), calculated using the getConvexHull function from the R library contoureR [55], was drawn around genomes from each clonal complex. Pairwise geographic distances between each isolate in kilometers were calculated using the distHaversine function from the R library geosphere [56]. The associations between genetic and spatial distance for clonal complexes Af2 and Eu3 were tested with a Mantel test (1000 permutations to assess significance) implemented using the R library ade4 [57].

### Putative transmission networks

Putative transmission networks based on a maximum inter-isolate pairwise SNP distance of 15 SNPs were defined using the R library iGRAPH [58] and plotted and annotated using the R library ggraph [59].

### Molecular dating of Eu3 clonal complex

BEAST v1.8.4 [60] was run on a SNP alignment of all Ethiopian (n = 40) and published (n = 67) Eu3 genomes (Supplementary File 2), using tip sampling dates for calibration. Three runs of 10^8^ Markov chain Monte Carlo (MCMC) iterations were performed using an HKY substitution model, strict or constant molecular clock and constant or exponential population size and growth (12 separate runs). The performance of each model was assessed through the comparison of posterior marginal likelihood estimates (MLE) and the model with the highest Bayes factor [61] (strict clock/constant population size) was selected (Supplementary Table 1). These three MCMC runs were combined using LogCombiner v1.8.4 (10% burnin) and convergence was assessed (posterior effective sample size (ESS) > 200 for each parameter) using Tracer v1.6. A maximum clade creditability tree summarizing the posterior sample of trees in the combined MCMC runs was produced using TreeAnnotator v1.8.4. The resulting tree was annotated using ggtree [62]. The R library TIPDATINGBEAST [63] was used to confirm a temporal signal in the dataset; briefly, the dates for each sample were resampled to generate 20 new datasets with randomly assigned dates. BEAST was then run on these new datasets using the same strict constant priors.

## Results

### Study sites and population

A total of 134 *M. bovis* isolates were successfully sequenced and passed QC; these included 127 isolates collected from 85 cattle, six isolates from six human TB patients and a single isolate from a dromedary. The origin of isolation and WGS based genotypes of these isolates are listed in Table 1 (the complete set of metadata can be found in Supplementary File 1). The majority of the isolates were collected from the large dairy belt in central Ethiopia, in the surroundings of Addis Ababa (Figure 1A). *In silico* spoligotype assignment revealed a total of 22 different spoligotypes with the most prevalent types being SB0134 (n = 36), SB1176 (n = 36) and SB0133 (n = 28). Three different clonal complexes were observed in the dataset (Figure 1B): African 2 (Af2; n = 92), European 3 (Eu3; n = 40) and Unknown8 (n = 2).

**Table 1.**
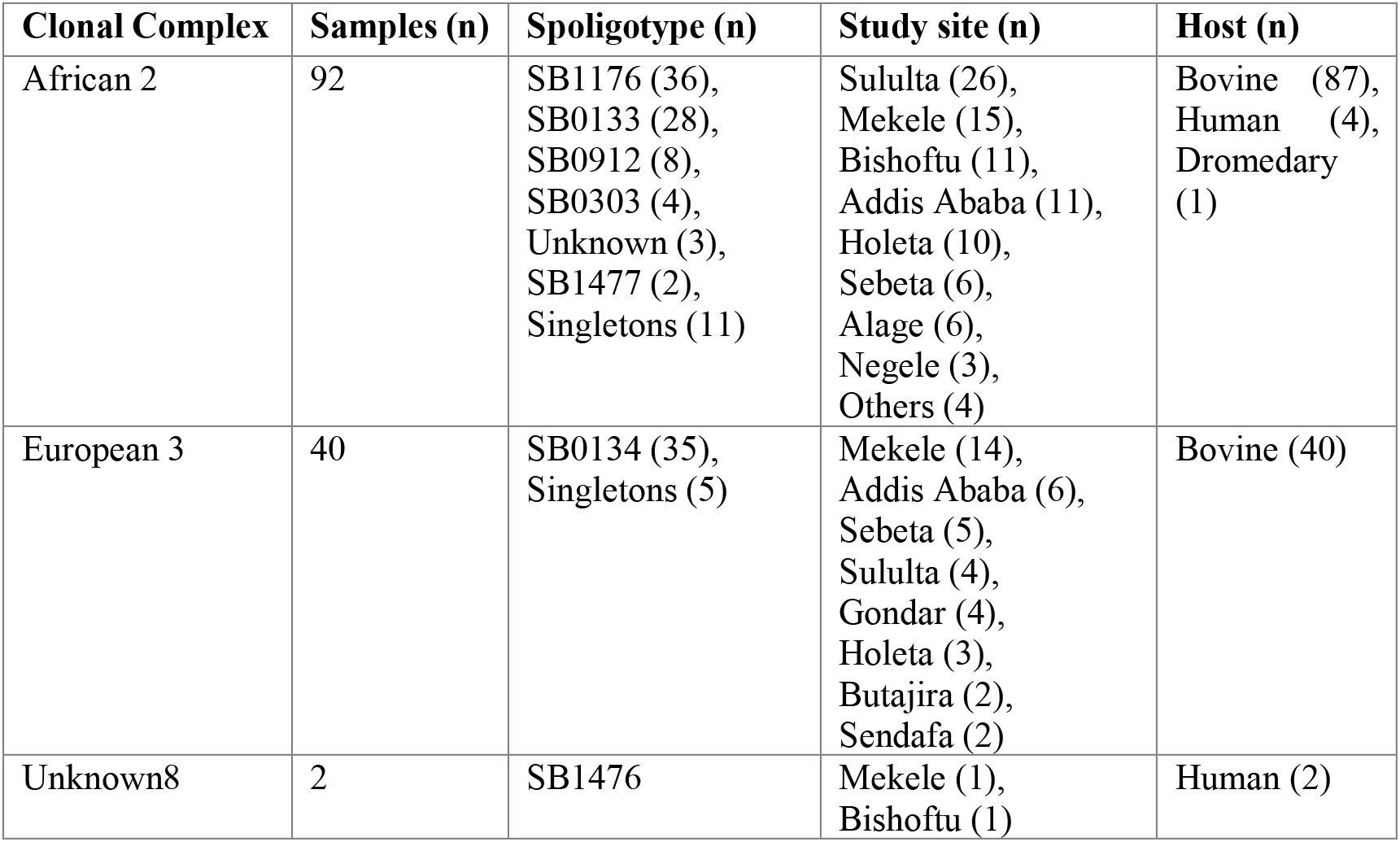
Metadata of 134 sequenced *Mycobacterium bovis* isolates included in this study, assigned to the African 2, the European 3, and the Unknown8 clonal complexes.

### Comparison of sequenced genotypes with previously published genotypes

Spoligotype information for previously published *M. bovis* isolates from animals, mainly cattle, was collated and the likely clonal complex inferred [64] (Table 2). The most frequent Af2 and Eu3 spoligotypes (SB1176, SB0133, SB0912, SB1477, and SB0134) previously identified in Ethiopia were also represented amongst the spoligotypes of the sequenced isolates in this study at similar frequencies. SB1476 (Unknown8), previously found exclusively in Ethiopian cattle, was found in two of the six isolates collected from humans.

**Table 2:** Genotyping data and inferred clonal complex of previously published *Mycobacterium bovis* isolates from Ethiopia.

| Clonal Complex | Samples (n) | Spoligotype (n) | Study site (n) | Host (n) | Reference (n) |
| --- | --- | --- | --- | --- | --- |
| African 2 | 190 | SB1176 (104),<br>SB0133 (50),<br>SB1477 (7),<br>SB0912 (6),<br>SB1490 (4),<br>SB0933 (3),<br>SB1491 (2),<br>SB1942 (2),<br>SB1983 (2),<br>Singletons (10) | Addis Ababa (64),<br>Holeta (41),<br>Kombolcha (26),<br>Negelle (24),<br>Jinka (12),<br>Adama (7),<br>Woldiya (4),<br>Gondar (3),<br>Fiche (3),<br>Ghimbi (2),<br>Butajira (2),<br>Melga-Wondo (1),<br>Hawassa (1) | Bovine (188),<br>Camel (2) | Berg 2009 [11](48),<br>Ameni 2007 [13](41),<br>Biffa 2010 [15](27),<br>Gumi 2012 [31](24),<br>Firdessa 2012 [8](19),<br>Ameni 2010 [12](13),<br>Tsegaye 2010 [18](7),<br>Mekibeb 2013 [17](6),<br>Ameni 2013 [14](3),<br>Mamo 2011 [19](2) |
| European 3 | 24 | SB0134 (19),<br>SB1517 (3),<br>Singletons (2) | Addis Ababa (21),<br>Yabello (2),<br>Gondar (1) | Bovine (24) | Firdessa 2012 [8](12), Biffa 2010 [15](6),<br>Berg 2009 [11](5),<br>Tsegaye 2010 [18](1) |
| Unknown8 | 18 | SB1476 | Ghimbi (8),<br>Gondar (7),<br>Butajira (2),<br>Jinka (1) | Bovine (18) | Berg 2009 [11](18) |

### Phylogenetic relationships of M. bovis genomes

The phylogenetic tree of 134 Ethiopian genomes, rooted with the Eu1 reference AF2122/97, shows a distinct phylogenetic structure with the three included clonal complexes clearly segregating in the phylogeny (Figure 1B). The Unknown8 clonal complex is basal followed by a split leading to Eu3 and Af2. Three distinct sub-lineages were observed for Af2 and each was associated with a predominant spoligotype (SB0303, SB0133 and SB1176).

### Spatial distribution of M. bovis clonal complexes

Figure 2A shows a spatial distribution of the 134 isolates overlaid with the smallest polygon incorporating the isolates from each clonal complex. There appears to be a distinct geographical distribution with Eu3 isolates more likely to be distributed to the north and west of the country whilst Af2 isolates have a more southern and eastern distribution. There is significant overlap between these two clonal complexes in central Ethiopia around Addis Ababa, likely reflecting a greater density of infected cattle from across the country.

**Figure 2.**
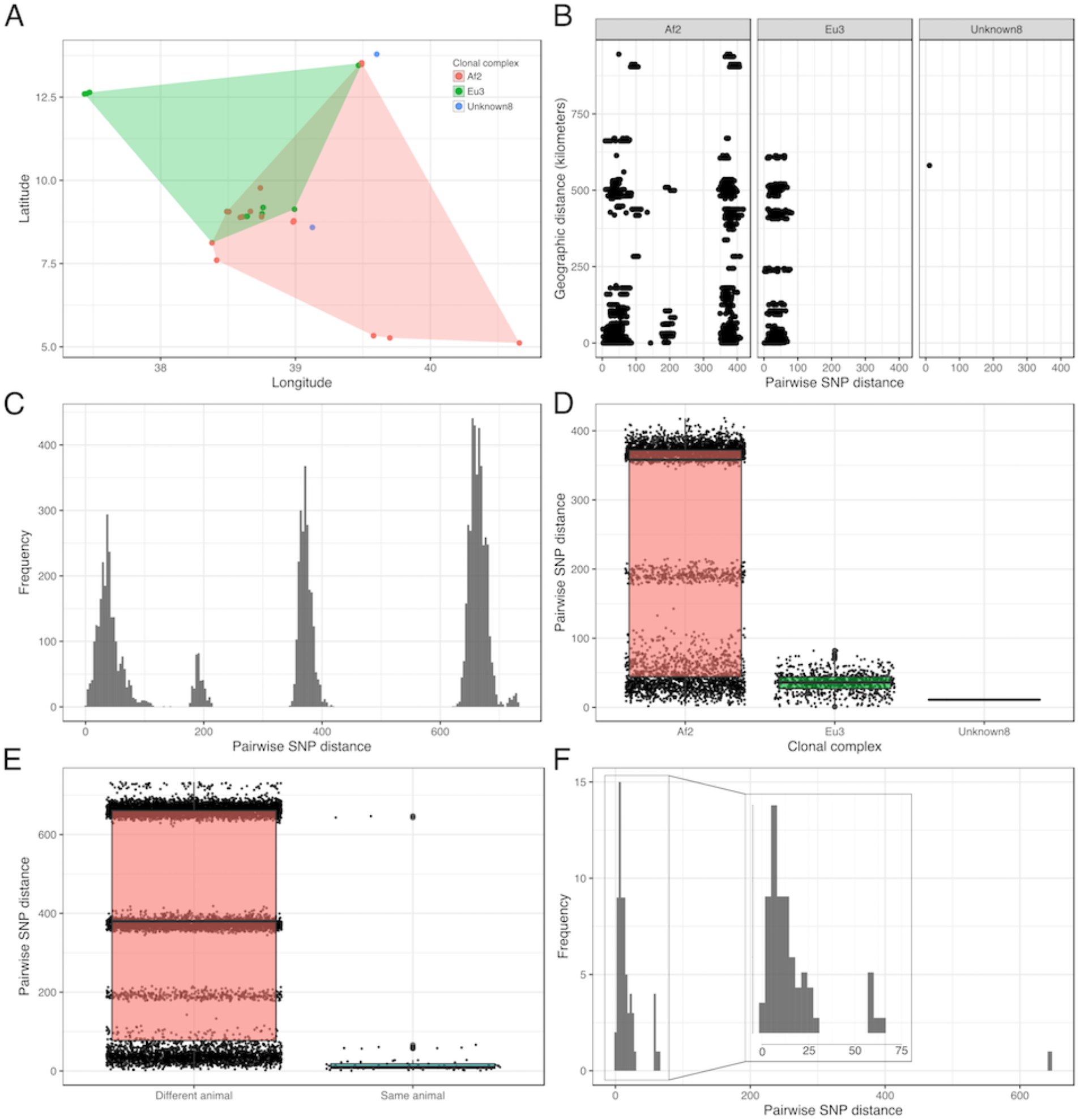
A. Spatial analysis of distribution of isolates coloured by clonal complex. Each polygon represents the minimum convex polygon of the sampled locations of the isolates from each clonal complex; B. Scatterplot of SNP distance against geographic distance for all pairs of genomes; C. Histogram of all pairwise SNP distances; D. Boxplot of all pairwise SNP distances separated by clonal complex; E. Boxplot of all pairwise SNP distances separated by within- and between-animal; F. Histogram of within-animal pairwise SNP distance. The insert shows the 0 – 75 SNP range zoomed in.

A comparison of pairwise genetic distance (SNPs) and geographic distance (kilometers) for all pairs of genomes is shown in Figure 2B. The results of the Mantel tests for Eu3 and Af2 (abattoir isolates removed) showed that there was no association between genetic and spatial distance (Eu3 observation = 0.10; simulated p-value: 0.001; Af2 observation = 0.03; simulated p-value: 0.001). Instead, the pattern observed reflects the large geographic distribution of these genetic lineages in Ethiopia.

### Genetic diversity and putative transmission

The distribution of pairwise SNP distances within the dataset was multimodal reflecting the population structure of the dataset with distinct modes observed for the three clonal complexes as well as the distinct sub-structure of Af2 (Figure 2C). Categorizing the pairwise SNP distances by clonal complex showed that there is considerably more diversity within Af2 compared to Eu3 and Unknown8 with the maximum pairwise SNP distance within Af2 being 418 SNPs, compared to 82 SNPs and 11 SNPs for Eu3 and Unknown8, respectively (Figure 2D). The within-host pairwise SNP diversity had a median of 10 SNPs (range 1 – 647 SNPs) whilst the between-host pairwise SNP diversity had a median of 380 SNPs (range 2 – 733 SNPs; Figure 2E). Further examination of the within-host diversity revealed three distinct peaks within the distribution of pairwise SNP distances with the majority of within-host isolates being within 1 and 30 SNPs of each other (Figure 2F). The maximum within-host pairwise SNP distance of 647 SNPs was observed in an animal (ser-002) with three isolates, two from Af2 and one from Eu3.

A total of 17 putative transmission clusters were defined using a pairwise SNP threshold of 15 SNPs. The networks varied in size between two and nineteen isolates, and there were 35 isolates that were not assigned to a network (singletons; n = 35; Figure 3B). Extensive within- and between-herd transmission was observed with both short- and long-distance transmission occurring. Examples of isolates from the same animal being in different transmission clusters were observed; for instance, isolates from animal ALB720 were found in two different transmission clusters along with isolates from animal 4589 (Figure 3B). These animals were from herds N (Alage) and Y (Mekele) which are approximately 662 kilometres apart. The largest transmission cluster, containing 19 isolates from 16 animals, showed clear epidemiological links between the Sululta abattoir (SA) and herds from around Sebeta (A, O, P, Q) and Bishoftu (S, T), whilst the second largest transmission cluster, containing 14 isolates from 11 animals showed links to abattoirs in and around Addis Ababa (AA, BA, SA) and herds from as far away as Gondar (U) and Mekele (X, Y).

**Figure 3.**
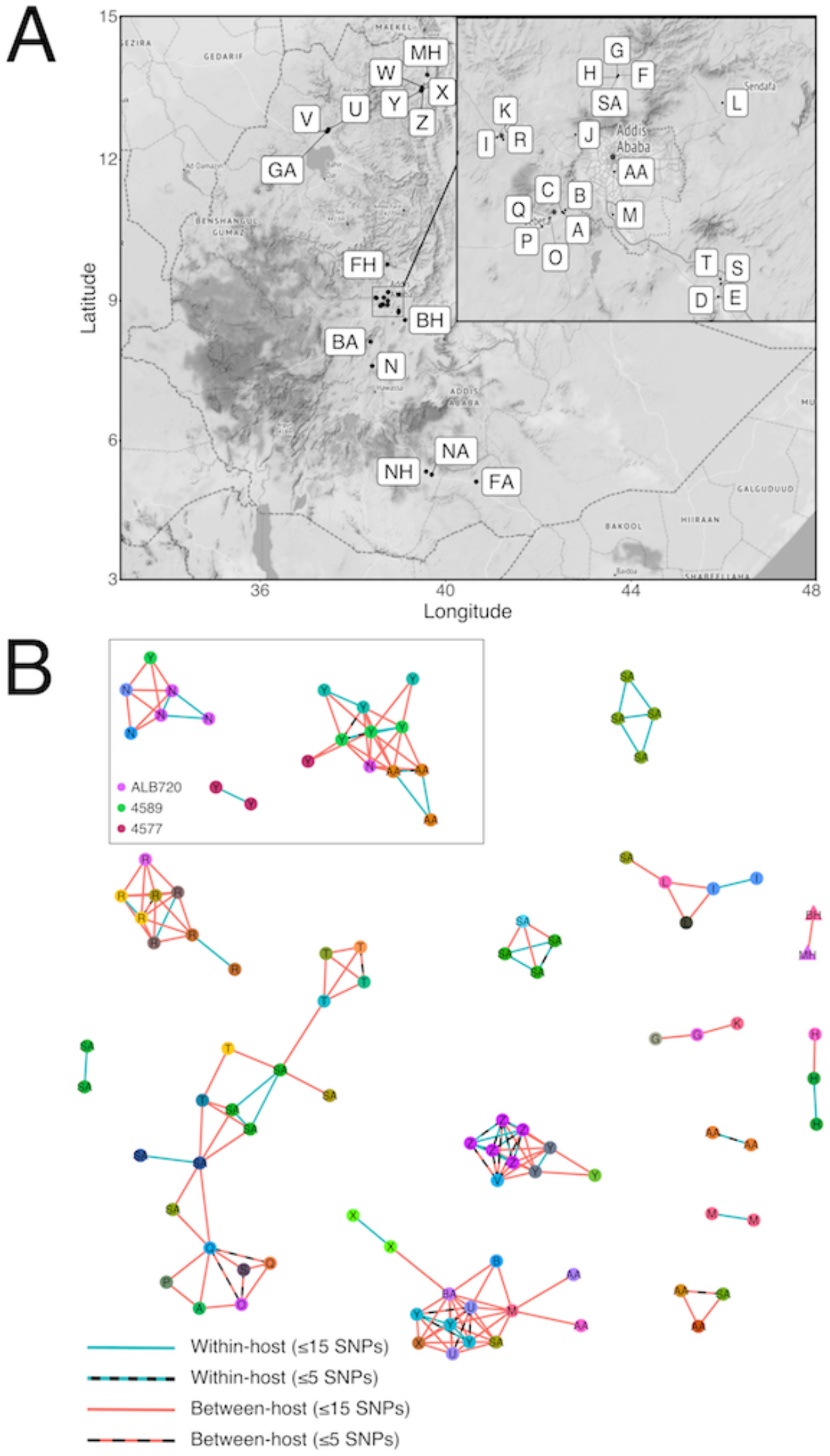
A. Map of Ethiopia showing the locations of the herds, abattoirs and hospitals from which the isolates were sourced (Letter coding from Supplementary File 1). The region around Addis Ababa was magnified in the insert; B. Putative transmission clusters defined using a pairwise SNP threshold of 15 SNPs. Nodes are coloured by animal and labelled with the herd, abattoir or hospital of isolation. Edges coloured in blue represent within-host links whilst edges coloured in blue represent between animal links. For simplicity, networks where n < 2 are not shown.

### Global and European 3 specific phylogenetics

Figure 4A shows a maximum likelihood phylogeny of 485 *M. bovis* genomes from 22 countries (including the genomes from this study), collected from 20 different host species between 1983 and 2018. All previously described clonal complexes along with one undescribed clonal complex (labelled as ‘Other’) are represented in the collection. The structure of the phylogeny is consistent with recently published work [40] and shows that pyrazinamide (PZA)-susceptible *M. bovis* from Malawi are basal to the other clonal complexes which are all PZA-resistant. The first of the next two ancestral splits in the phylogeny leads to the sister clades Af2 and Eu3, where the majority (n = 132) of the Ethiopian genomes are contained, and the clade containing all other clonal complexes (Other, Af1, Eu1, Eu2 and Unknown3-8). The remaining two Ethiopian genomes (SB1476/Unknown8) are the outgroup to the globally distributed Eu1 clonal complex (Figure 4A).

**Figure 4.**
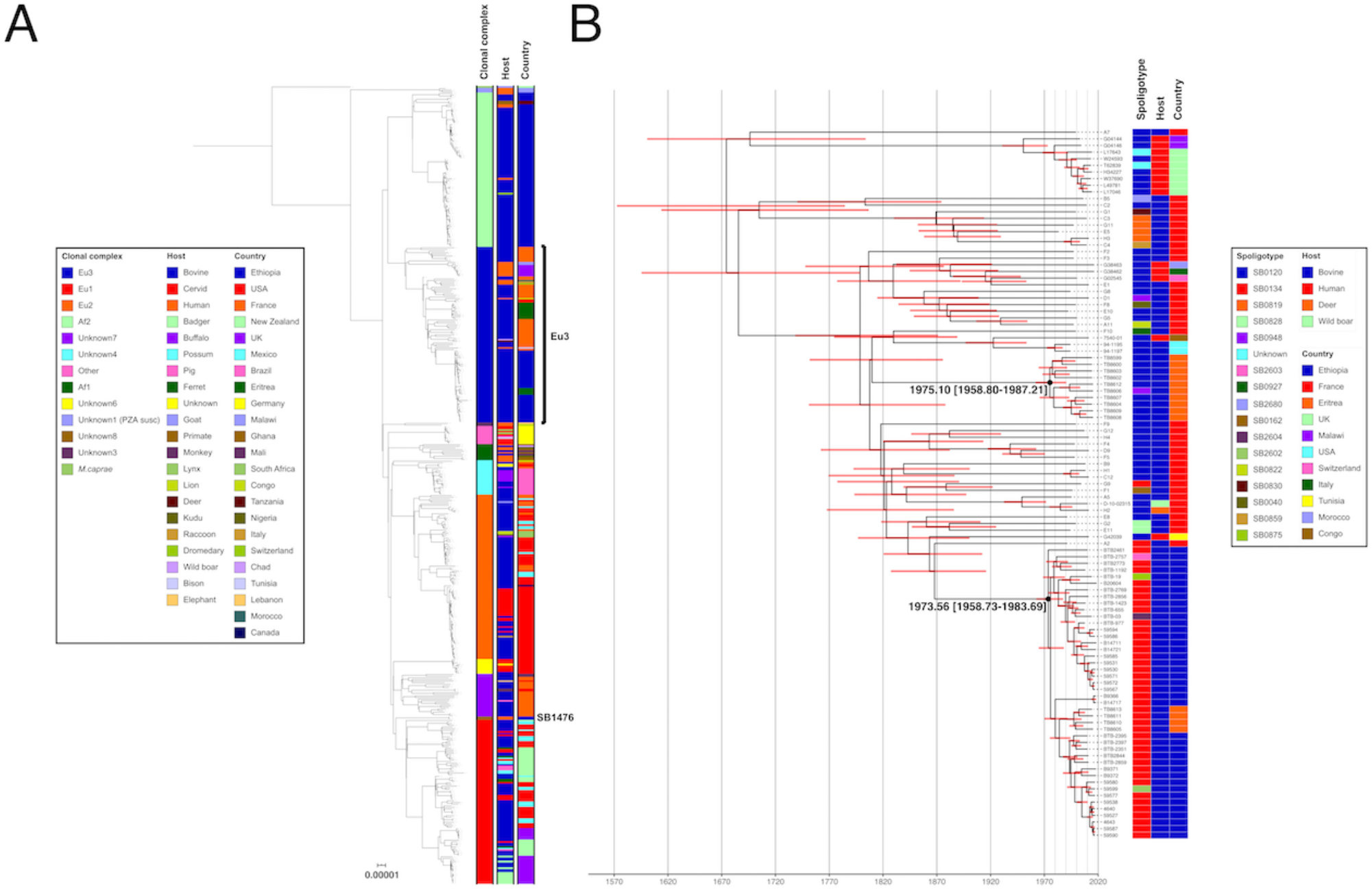
A. Global maximum likelihood phylogeny of 485 *Mycobacterium bovis* genomes rooted using *Mycobacterium caprae;* B. Time-calibrated maximum clade credibility tree of European 3 genomes.

The 40 Ethiopian Eu3 genomes were combined with 67 other Eu3 genomes from 10 countries and the resulting phylogeny was dated using BEAST (Figure 4B). The median date of the most recent common ancestor (MRCA) for this clade was estimated to be 1674 [95% CI: 1557-1769]. The dated phylogeny shows two potential Eu3 introduction events into Ethiopia (Eritrea was part of Ethiopia until 1993) around the same time period: 1975 [95% CI: 1959-1987] and 1974 [95% CI: 1959-1984] (Figure 4B). The Eu3 phylogeny also contains nine BCG vaccine isolates collected in Malawi and the UK and these have a MRCA dating back to 1950 [95% CI: 1926-1968]. The results of the dated tip randomization analysis are shown in Supplementary Figure 1. The substitution rate in the observed data did not overlap with the estimated substitution rates in the randomized datasets showing that the temporal signal observed was not obtained by chance.

## Discussion

This is the first study to use WGS to examine the population structure of *M. bovis* in Ethiopia. A total of 134 isolates from 12 study sites across Ethiopia were successfully sequenced and three distinct clonal complexes, which appeared to be mostly geographically segregated, and 22 different spoligotypes were observed in the dataset. The in-depth genome analysis showed there was no association between geographic and genetic distance and differing levels of genetic diversity were observed amongst the three clonal complexes. A range of within-host diversity was also observed with isolates from the same animal often being in different transmission clusters, as well as evidence of both short- and long-distance transmission with isolates in transmission clusters collected hundreds of kilometres apart. Molecular dating of a collection of Eu3 genomes which included the Ethiopian genomes in this study estimated the date of introduction of different Eu3 sub-lineages into Ethiopia and Eritrea during the same time period during the early 1970s (CI: 1958-1987).

To get a better understanding of how well the genomes analysed in this study represented the known population structure of *M. bovis* in Ethiopia, we collated spoligotype information for the vast majority of Ethiopian *M. bovis* isolates from animals (mainly cattle) published to date [64]. When comparing the spoligotypes in the current study (Table 1) with those previously published (Table 2), it was clear that the most frequent spoligotypes of Af2 and Eu3 identified in Ethiopia (SB1176, SB0133, SB0912, SB1477, and SB0134) were also represented in our dataset with their prevalence observed at a similar frequency. Spoligotype SB1476 has also been frequently observed in cattle in previous studies [11] but we provide the first evidence of this spoligotype in humans, possibly as a result of zoonotic transmission. On this basis, the isolates sequenced in this study appear to be representative of the population structure of *M. bovis* in Ethiopia.

Examination of the geographical distribution of the clonal complexes, in particular Af2 and Eu3, contained within the dataset showed a potential geographic divide between their distributions with an overlap in the herds around Addis Ababa. The number of isolates from outside this region is small but given the expected movement, often long-distance, of animals between different herds, the geographic separation of clonal complexes is still distinct. The reason for this pattern is unclear but may be due to historical isolation of different parts of the country due to geographic distance; it is interesting that Af2, the indigenous lineage in Ethiopia, was not found in the north west of the country but this may simply be due to a lack of sampling. We could find no evidence of a relationship between genetic and geographic distance for the different clonal complexes implying that the *M. bovis* population in Ethiopia is well-mixed and maintained by both short- and long-distance cattle movements. There was a large difference observed in genetic diversity between clonal complexes Af2 and Eu3; this is likely due to the long-term endemic nature of Af2 in Ethiopia which has allowed for significant genetic divergence over time with the emergence of clear sub-lineages within this clonal complex observed. Conversely, the comparatively recent introduction of Eu3 into Ethiopia (see below), has not allowed enough time for more genetic diversity to emerge. As expected, considerable genetic diversity was observed between isolates from different animals with some isolates, from different clonal complexes, being as much as 733 SNPs apart. Less expected was the range of within-host diversity observed. The majority of isolates from the same animal were up to 30 SNPs (median = 10 SNPs) apart from each other; given the previously estimated mutation rate of *M. bovis* of 0.15 – 0.53 SNPs per genome per year [65, 66] it is likely infections are being maintained over a long period of time. Less closely related isolates from the same animal were also found showing that multiple infections by different strains was taking place amongst our samples.

With bTB being endemic in Ethiopia and the intensive dairy sector highly affected, there is considerable interest from stakeholders to explore control strategies for this disease. One of the aims of the ETHICOBOTS project was to use WGS to identify potential transmission of bTB in Ethiopian dairy cattle. We defined transmission clusters (Figure 3B) using a conservative threshold of 15 SNPs which would allow for both the identification of older transmission events but also allow for varying rates of mutation within the animals sampled. Using this threshold, we could find clear evidence of transmission occurring between animals from the same herd as well as animals from different herds and animals sampled at local abattoirs. We also found evidence of isolates from the same animal in different transmission clusters, showing that animals are being re-infected with different strains. There was also strong evidence of both short- and long-distance transmission with isolates in some transmission clusters being hundreds of kilometres apart. These findings are not surprising as very few farms in Ethiopia have the ability to control bTB in their herds, *e.g*. by test-and-slaughter based on the tuberculin skin-test and there is considerable long-distance trade between affected herds. Instead, this uncontrolled chronic disease results in cattle infected with *M. bovis* being likely to survive for a long enough time to be subject to exposure to multiple bTB strains and/or cattle trade (short- or long-distance), increasing the risk of disease transmission and reinfection.

There is considerable heterogeneity with respect to the size of defined transmission clusters and contribution from individual farms. Genomes isolated from Farm Y (Figure 3) are part of three clusters suggesting epidemiological links both locally and nationally. Although we must be cautious in interpreting patterns within such a sparse sample, this pattern is consistent with so-called “super-spreading” behaviour where a small number of herds contribute disproportionately to transmission. The potential for such super-spreading, with an opportunity to target herds for control, was evident in our movement-based network analysis [67].

Additionally, we wanted to examine the possibility of zoonotic transmission between humans and cattle in Ethiopia. The minimum pairwise SNP distance between any pairs of human and animal isolates was 41 SNPs (median 658 SNPs) providing no evidence of recent zoonotic transmission from the cattle or herds that were sampled (potential transmission events would be highlighted by small pairwise SNP distances). This was unsurprising, given there were only six isolates from humans included in this study. However, two of the human isolates (of type SB1476) were only 11 SNPs apart from each other suggesting a potential epidemiological link; these isolates were from Bishoftu and Mekele, approximately 581 kilometres apart, suggestive of long-distance travel of infected individuals or animals. Overall, these data provide clear evidence of considerable transmission between cattle but denser sampling will be required to establish specific evidence of zoonotic transmission.

Recent work has suggested an East African origin for *M. bovis* [40]. Given its position in the global phylogeny and confined geographical distribution [20], Af2 has likely been circulating in East Africa for hundreds of years; unfortunately, the lack of genomes from other parts of the world to provide context to the Ethiopian Af2 genomes, due to the small number sequenced, means confirming this in Ethiopia is currently not possible. However, we can show that there is a clear phylogenetic structure in this clonal complex with the Ethiopian genomes dividing into three distinct lineages. The position of Eu3 as the sister group to Af2 also implies a likely East African origin; however, based on the samples available for this study, the basal position of French isolates (Figure 4B) suggests that the currently circulating Eu3 lineage in Ethiopia may in fact be European in origin. One possible explanation for this is that the ancestor of Eu3 was brought to Europe a few hundred years ago and that the modern Ethiopian and Eritrean genomes are descendants of that population, not of the ancestral population that may or may not still be circulating somewhere in East Africa. What is reasonably clear, given the long branches and subsequent expansion, is that there were two introductions of Eu3 sub-lineages, consistent with the study that first analysed the Eritrean WGS data [25], into Ethiopia between 1958 and 1987 (which also included Eritrea during that time) with the median estimate for introduction being in the early 1970s. Given the very similar dates, it is possible that these were part of the same series of cattle imports. In terms of likely origin of these imports, France should be viewed as a proxy for the *M. bovis* diversity seen in mainland Europe (these samples were chosen to represent the diversity seen in France [47]), so the actual origin may be elsewhere in Europe. There are historical records of the first dairy cattle being imported into Ethiopia around 1950 as part of the United Nations Relief and Rehabilitation Administration (UNRRA) [68] with further subsequent imports from Kenya in 1959 [69]. Several livestock and dairy development projects that took place in the 1950-70s and funded by Sweden and the World Bank may have brought in dairy cattle of exotic breeds from overseas [3].

The other clonal complex found in Ethiopia, Unknown8, represented by two human genomes with the spoligotype SB1476, has thus far only been found in Ethiopia [11]. The position of this lineage in the global tree, and the hypothesized East African origin of *M. bovis* [40], suggests that this clonal complex may be the ancestor of Eu1, the most prevalent and geographically distributed *M. bovis* lineage known to date. However, further work would need to be done to confirm this through the collection of larger numbers of isolates with spoligotype SB1476.

This study performed the first WGS-based analysis of *M. bovis* isolates from Ethiopia and attempted to include isolates from multiple herds in different parts of the country. There is considerable genetic diversity amongst *M. bovis* in Ethiopia with multiple clonal complexes circulating and that they were likely to have been introduced in the country at different time-points. This work is important as it helps to better understand bTB transmission in cattle in Ethiopia and will potentially inform national strategies for bTB control in Ethiopia and beyond.

## Authors and contributors

Conceptualization: James LN Wood, Stefan Berg, Andrew JK Conlan, R Glyn Hewinson, Gobena Ameni. Data curation: Stefan Berg, Andries J van Tonder, Getnet Abie Mekonnen, Gizat Almaw. Formal analysis: Stefan Berg, Andries J van Tonder, Eleftheria Palkopoulou, Javier Nunez-Garcia. Investigation: Gizat Almaw, Getnet Abie Mekonnen, Hawult Taye, Aboma Zewude, Mekedes Tamiru, Abebe Olani, Abde Aliy, Matios Lakew, Melaku Sombo, Colette Diguimbaye, Markus Hilty, Adama Fané, Borna Müller. Methodology: Gizat Almaw, Getnet Abie Mekonnen, James LN Wood, Stefan Berg, Andries J van Tonder, Richard J Ellis. Project administration: Solomon Gebre, Adane Mihret, Stefan Berg, James LN Wood. Supervision: Adane Mihret, Tamrat Abebe, Gobena Ameni, Solomon Gebre, Stefan Berg, Andries J van Tonder. Writing – original draft: Stefan Berg, Andries J van Tonder, Getnet Abie Mekonnen, Gizat Almaw. Writing – review & editing: Andrew JK Conlan, Adane Mihret, Tamrat Abebe, Gobena Ameni, Balako Gumi, Julian Parkhill, James LN Wood, Eleftheria Palkopoulou, Richard J Ellis, R Glyn Hewinson

## Ethical approval

Ethical approvals to implement the research of ETHICOBOTS were granted by the National Research Ethics Review Committee (NRERC No. 3.10/800/07), and by the institutional review boards at Aklilu Lemma Institute of Pathobiology, Addis Ababa University (Reference number IRB/ALIPB/2018), and at AHRI-ALERT (Proj. Reg. no P046/14). Enrolment of human study participants was done after written informed consent was secured and signed agreements taken from all eligible participants.

## Acknowledgements

The members of the ETHICOBOTS consortium are: Abraham Aseffa, Adane Mihret, Bamlak Tessema, Bizuneh Belachew, Eshcolewyene Fekadu, Fantanesh Melese, Gizachew Gemechu, Hawult Taye, Rea Tschopp, Shewit Haile, Sosina Ayalew, Tsegaye Hailu, all from Armauer Hansen Research Institute, Ethiopia; Rea Tschopp from Swiss Tropical and Public Health Institute, Switzerland; Adam Bekele, Chilot Yirga, Mulualem Ambaw, Tadele Mamo, Tesfaye Solomon, all from Ethiopian Institute of Agricultural Research, Ethiopia; Tilaye Teklewold from Amhara Regional Agricultural Research Institute, Ethiopia; Solomon Gebre, Getachew Gari, Mesfin Sahle, Abde Aliy, Abebe Olani, Asegedech Sirak, Gizat Almaw, Getnet Mekonnen, Mekdes Tamiru, Sintayehu Guta, all from National Animal Health Diagnostic and Investigation Center, Ethiopia; James Wood, Andrew Conlan, Alan Clarke, all from Cambridge University, United Kingdom; Henrietta L. Moore and Catherine Hodge, both from University College London, United Kingdom; Constance Smith at University of Manchester, United Kingdom; R. Glyn Hewinson from Aberystwyth University, United Kingdom; Stefan Berg, Martin Vordermeier, Javier Nunez-Garcia, all from Animal and Plant Health Agency, United Kingdom; Gobena Ameni, Berecha Bayissa, Aboma Zewude, Adane Worku, Lemma Terfassa, Mahlet Chanyalew, Temesgen Mohammed, Miserach Zeleke, all from Addis Ababa University, Ethiopia. We thank the Ethiopian Ministry of Agriculture and the Ethiopian Ministry of Health for their support to the ETHICOBOTS project.

## Supplementary Tables and Figures

**Supplementary Table 1:** Model performance based on marginal likelihood estimates (MLE) Bayes factors.

| Model | log MLE | log Bayes Factor | Strength of Evidence (Kass & Raftery, 1995) |
| --- | --- | --- | --- |
| Relaxed constant | -5604954.493 | - | - |
| Relaxed exponential | -5604956.354 | 1.861248229 | Positive |
| Strict exponential | -5605103.905 | 149.4115816 | Very strong |
| Strict constant | -5605106.444 | 151.9514509 | Very strong |

**Supplementary Figure 1:**
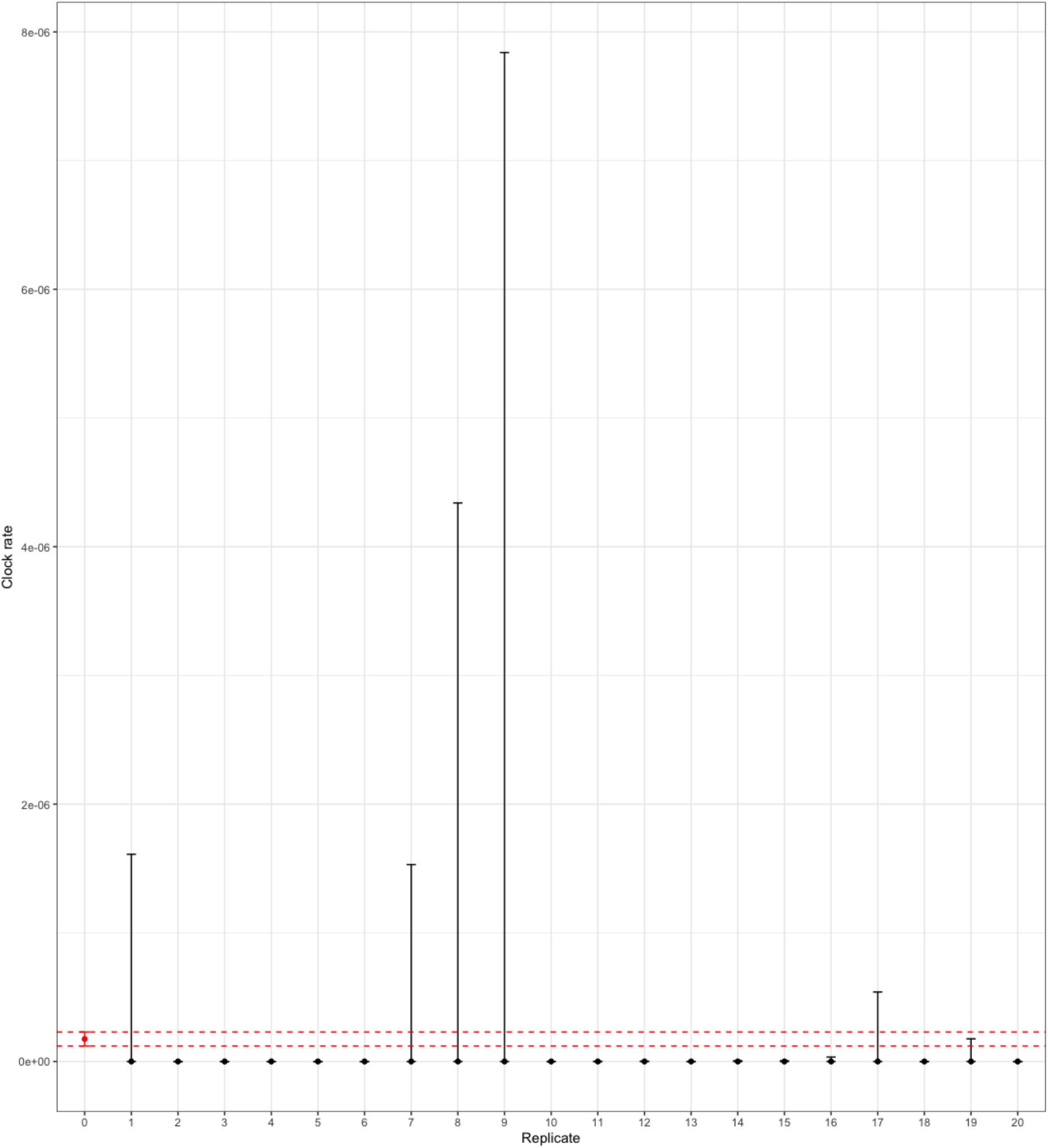
BEAST dated tip randomization (DTR) analysis. Observed data is highlighted in red.

